# Causal dissociation of occipital and mid-fusiform contributions to human visual experience

**DOI:** 10.64898/2026.11.25.769180

**Authors:** Dounia Hajhajate, Valerio Frazzini, Sergio Della Sala, Alessandro Cocuzza, Jianghao Liu, Katia Lehongre, Sara Fernandez Vidal, Vincent Navarro, Paolo Bartolomeo

**Affiliations:** Sorbonne Université, Institut du Cerveau - Paris Brain Institute - ICM, Inserm, CNRS, APHP, Pitié-Salpêtrière Hospital, 47 Bd de l’Hôpital, 75013 Paris, France; Epilepsy and EEG Unit, Département de Neurologie and Reference Center for Rare Epilepsies, ERN Epicare, AP-HP, Pitié-Salpêtrière Hospital, 47 Bd de l’Hôpital, 75013 Paris, France; Human Cognitive Neuroscience, Department of Psychology, University of Edinburgh, UK; ICoN Cognitive Neuroscience Center, University School for Advanced Studies, IUSS, Pavia, Italy

## Abstract

Visual experiences can arise without corresponding retinal input, as in imagery, dreams, and hallucinations. Influential models of visual mental imagery propose that mental images acquire their visual quality in early visual cortex. Direct intracranial electrical cortical stimulation (DIECS) causally probes which cortical sites can evoke visual experiences, and of what kind. We examined de novo visual experiences evoked by stimulation in three patients undergoing stereo-EEG monitoring for drug-resistant epilepsy and localized the effective contact pairs using the Jülich-Brain cytoarchitectonic atlas. Among the sampled sites, stimulation of occipital and adjacent intraparietal cortex and of the posterior fusiform area FG1 elicited simple or intermediate phenomena, including flashes, colored spots, and elementary shapes. By contrast, only stimulation of sites in or adjacent to the mid-fusiform areas FG3 and FG4 elicited complex images of people, scenes, and objects. These findings provide cytoarchitectonically localized causal evidence that stimulation of right mid-fusiform cortex can trigger complex visual experiences, possibly through distributed networks. They also identify the mid-fusiform gyrus as a promising causal node for complex visual content, and are compatible with lesion and neuroimaging evidence challenging the necessity of early visual cortex for visual imagery.

## Introduction

Visual experiences arise during perception, mental imagery, dreams, and hallucinations. The cortical mechanisms supporting these experiences, and particularly the contribution of early visual cortex, remain debated. Influential accounts propose that visual imagery reinstates perceptual representations in early visual areas [1], whereas neuropsychological evidence challenges the necessity of these areas for visual imagery [2].

Direct intracranial electrical cortical stimulation (DIECS) allows investigators to identify cortical sites from which visual phenomena can be evoked, although stimulation effects may spread beyond the stimulated contacts. In their classic summary of 1,288 operations in 1,132 patients, with stimulation of the exposed cortical surface, Penfield and Perot [3] found that occipital stimulation predominantly elicited elementary visual phenomena, such as phosphenes. In contrast, temporal stimulation, particularly of the right hemisphere, could evoke complex images of people, scenes, and objects. In a subsequent study using subdural electrodes, Lee et al. [4] classified evoked visual phenomena as simple (spots and flashes), intermediate (geometric shapes), or complex (recognizable objects and scenes), and described a broadly posterior-to-anterior progression across these classes. However, surface electrodes offer limited access to the sulcal and basal cortex, making it difficult to precisely attribute these responses to specific anatomical regions.

Combining functional imaging with intracranial stimulation can relate evoked percepts to the visual selectivity of the stimulated cortex [5]. Using stereo-electroencephalography (sEEG), which provides access to sulcal and deep cortical regions, Jonas et al. [6] studied 22 patients and found that the proportion of stimulation sites eliciting visual phenomena (hallucinations, illusions, or recognition impairments) decreased from the occipital pole to the most anterior temporal sites sampled, with a more pronounced decline in the left hemisphere. Complex hallucinations consisted of faces and were evoked at seven right-hemisphere sites: five in the cuneus/parieto-occipital sulcus, one in the precuneus, and one in the fusiform gyrus. In a separate intracranial recording study of 140 patients, Jacques et al. [7] found increasing face selectivity along the ventral occipito-temporal cortex, with broadly similar onset latencies of face-selective activity across regions. This finding challenges a strictly serial model of ventral-stream processing but does not establish which sites can evoke complex visual experiences when stimulated. In particular, the role of the fusiform gyrus and its subdivisions remains uncertain.

We examined the visual experiences elicited by DIECS in three patients undergoing sEEG monitoring, with electrodes sampling lateral, basal, and medial occipito-temporal regions. We assigned effective stimulation sites to cytoarchitectonic areas using the Jülich-Brain atlas [8], focusing on subdivisions of the fusiform gyrus. This approach allowed us to relate the phenomenology of stimulation-evoked visual experiences to more precisely defined occipital and temporal sites. Complex visual experiences arose only from right mid-fusiform sites in or adjacent to the cytoarchitectonic areas FG3 and FG4 [9], whereas the sampled occipital sites and the posterior fusiform area FG1 [10] elicited only simple or intermediate phenomena.

## Methods

Data were obtained from three patients admitted to the Epilepsy Unit of the Department of Neurology, Pitié-Salpêtrière Hospital, Paris, France. All patients underwent sEEG monitoring as part of their presurgical evaluation for focal drug-resistant epilepsy. DIECS was performed during sEEG monitoring to map cortical function and delineate the epileptogenic zone. Patients were included if (1) electrodes had been implanted in the occipito-temporal cortex and (2) stimulation elicited visual experiences from at least two of the three classes defined by Lee et al. [4].

### Participants

Patient 1 was a 37-year-old left-handed man whose seizures began at 4 years of age. He had focal impaired awareness seizures that occasionally progressed to bilateral tonic-clonic seizures. At the time of investigation, he was treated with carbamazepine (800 mg/day), levetiracetam (2,500 mg/day), and zonisamide (200 mg/day). Brain MRI revealed subcortical band heterotopia involving the parieto-temporal regions. Bilateral sEEG exploration used 13 electrodes targeting the posterior quadrant of each hemisphere, sampling the parieto-occipital cortices, posterior lateral temporal cortices, and posterior right hippocampus. The epileptogenic zone was localized to the right lateral occipital cortex.

Patient 2 was a 26-year-old right-handed man whose seizures began at 5 years of age. Seizures occurred predominantly during sleep and started with a dream-like feeling of strangeness, followed by impaired awareness and bilateral upper-limb motor manifestations. At the time of investigation, he was treated with topiramate (300 mg/day) and valproic acid (1,000 mg/day). Brain MRI showed a large cortical lesion of the right lateral occipito-temporal region associated with periventricular nodular heterotopia. Nine electrodes sampled the lateral and medial right temporal cortex and the first and second right occipital gyri. The epileptogenic zone was localized to the posterior right temporal cortex.

Patient 3 was a 25-year-old right-handed woman who presented with drug-resistant focal epilepsy beginning in adolescence. Her seizures were characterized predominantly by visual and cognitive symptoms. Seizure onset typically involved complex visual hallucinations of moving objects or scenes, such as shelves, balloons, or trains, accompanied by dizziness or vertigo and experiential phenomena, including déjà vu and déjà vécu. These initial symptoms were followed by impaired awareness and complex motor automatisms, notably stereotyped towel-folding movements. At the time of presurgical evaluation, she was receiving eslicarbazepine (800 mg/day) and lacosamide (200 mg/day). Structural brain MRI was unremarkable, whereas PET-MRI revealed mild bilateral mesial temporal hypometabolism. The patient underwent two sequential sEEG explorations to delineate the epileptogenic zone. During the first exploration, 11 intracerebral electrodes were implanted in the right hemisphere, sampling the temporal pole, basal temporal cortex, hippocampus, amygdala, temporo-occipital junction, and second and third occipital gyri. Ictal sEEG recordings suggested initial involvement of the posterior basal temporal region, with rapid propagation to mesial temporal structures. However, the precise epileptogenic zone remained insufficiently defined, prompting a second sEEG exploration with eight intracerebral electrodes targeting the right posterior basal temporal and occipital cortices. The second sEEG investigation permitted a more precise delineation of the epileptogenic network, identifying the posterior right fusiform gyrus and the lateral temporo-occipital sulcus as key regions involved in seizure generation.

### sEEG recordings

Each electrode (Ad-Tech®, Oak Creek, WI, USA) comprised 4–12 platinum contacts, each 2.41 mm in length and 1.10 mm in diameter. Clinical requirements and the hypothesized location of the epileptogenic zone determined the number and trajectories of the electrodes, their anatomical targets, and the number of contacts. This hypothesis rested on each patient’s seizure semiology, scalp EEG, structural MRI, and, where available, metabolic imaging.

### DIECS protocol

DIECS was performed at the bedside in a quiet environment during 1–4-h sessions, conducted once or twice daily according to clinical judgment and patient fatigue. Patients were informed that stimulation would occur but not when, and were instructed to report any subjective experience spontaneously. When a response occurred, open-ended questions followed by targeted prompts were used to characterize its phenomenology, onset, offset, duration, and intensity, the temporal sequence of multiple manifestations, and their possible correspondence to habitual seizure symptoms.

The electrical stimulations retained in the primary analysis consisted of biphasic square-wave pulses delivered between adjacent contacts in a bipolar configuration at 50 Hz, with a pulse width of 0.5 ms per phase and a train duration of 5 s [11]. Bipolar stimulation is relatively focal, although its spatial extent depends on electrode geometry, tissue conductivity, and current intensity [12]. We adjusted stimulation intensity for each contact pair based on the cortical site, the patient’s epilepsy profile, and the responses elicited. Intensities ranged from 1 to 4 mA in the primary analysis (Table 1) and from 0.5 to 8 mA in the additional trials (Table S1), which also included stimulation at 1 Hz.

**Table 1:**
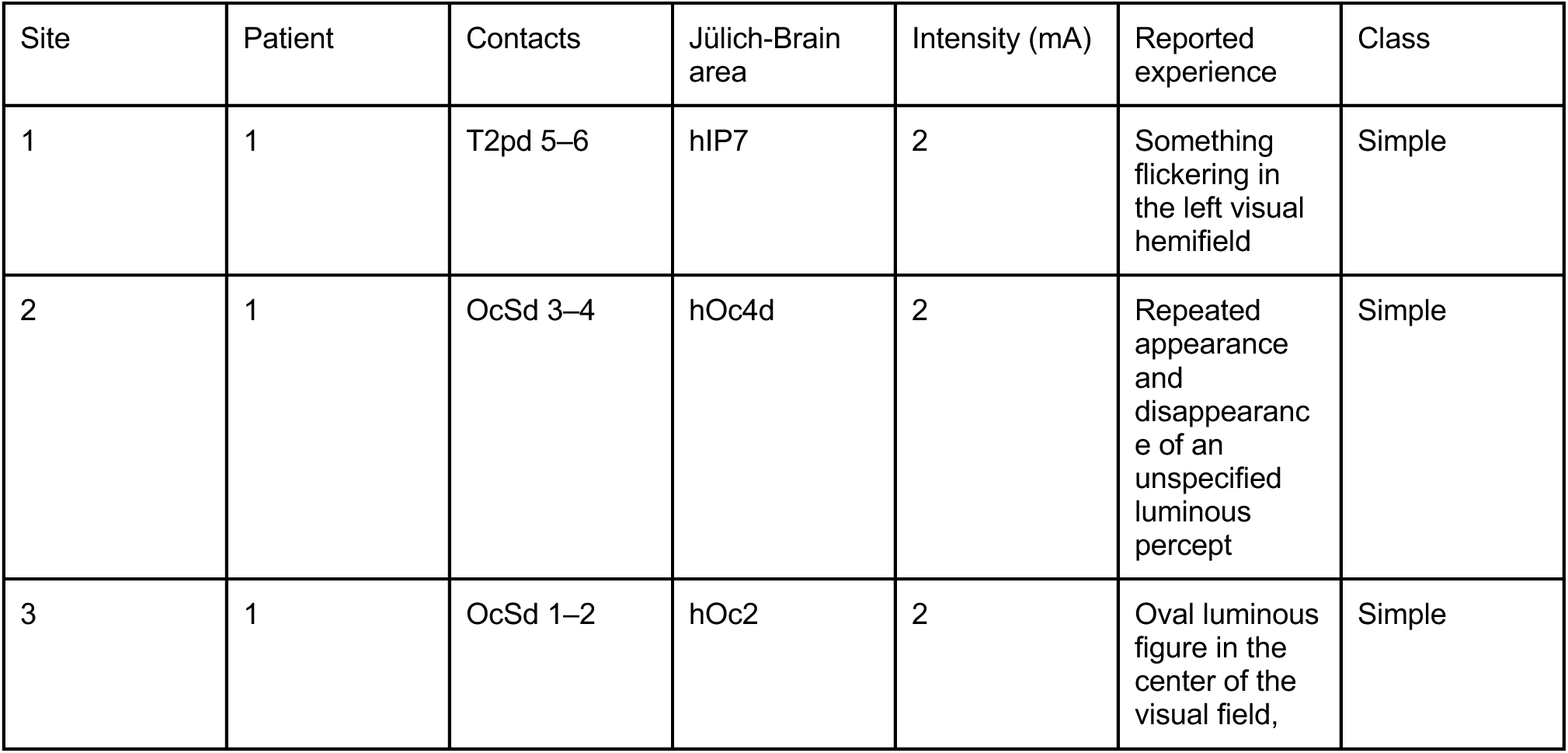
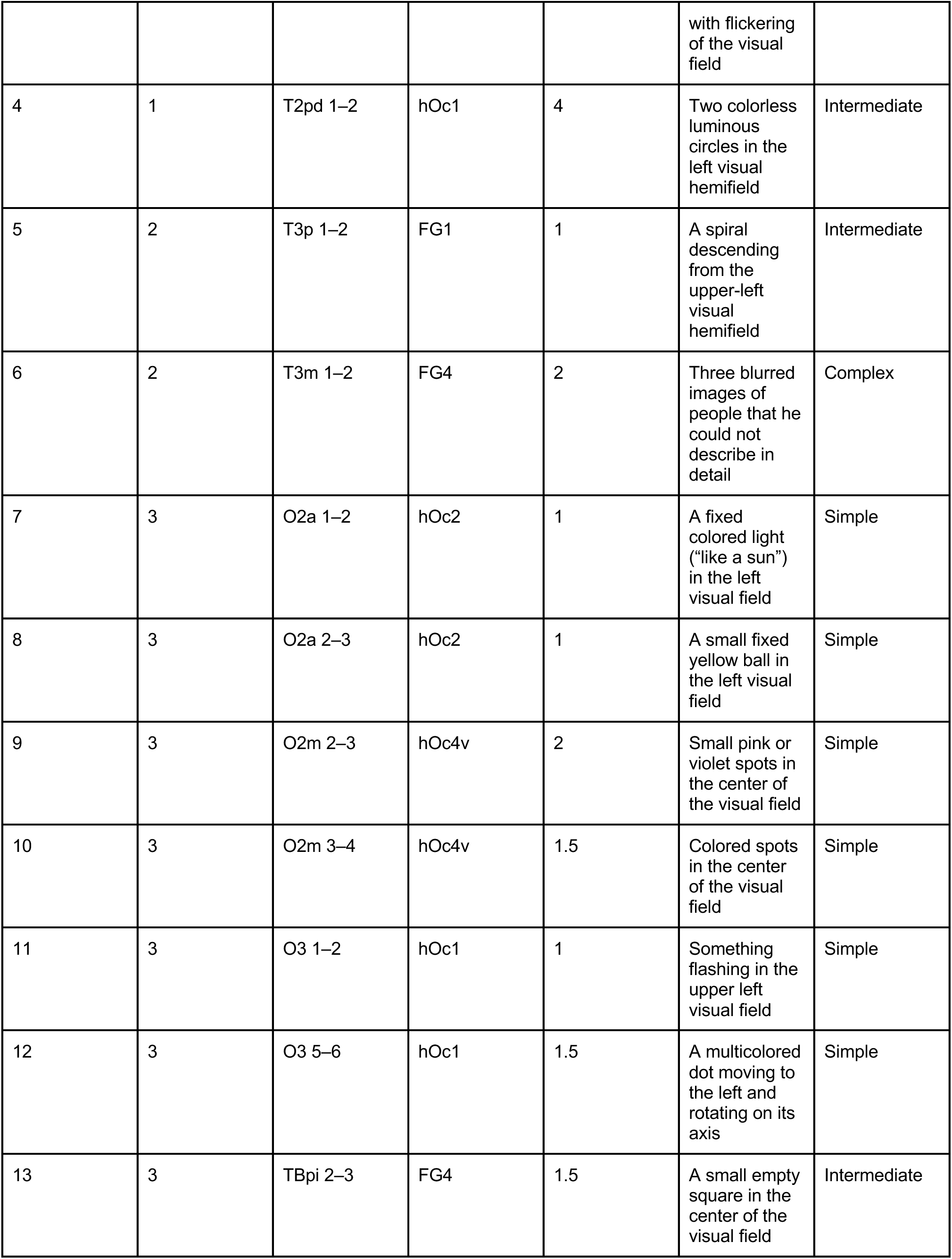
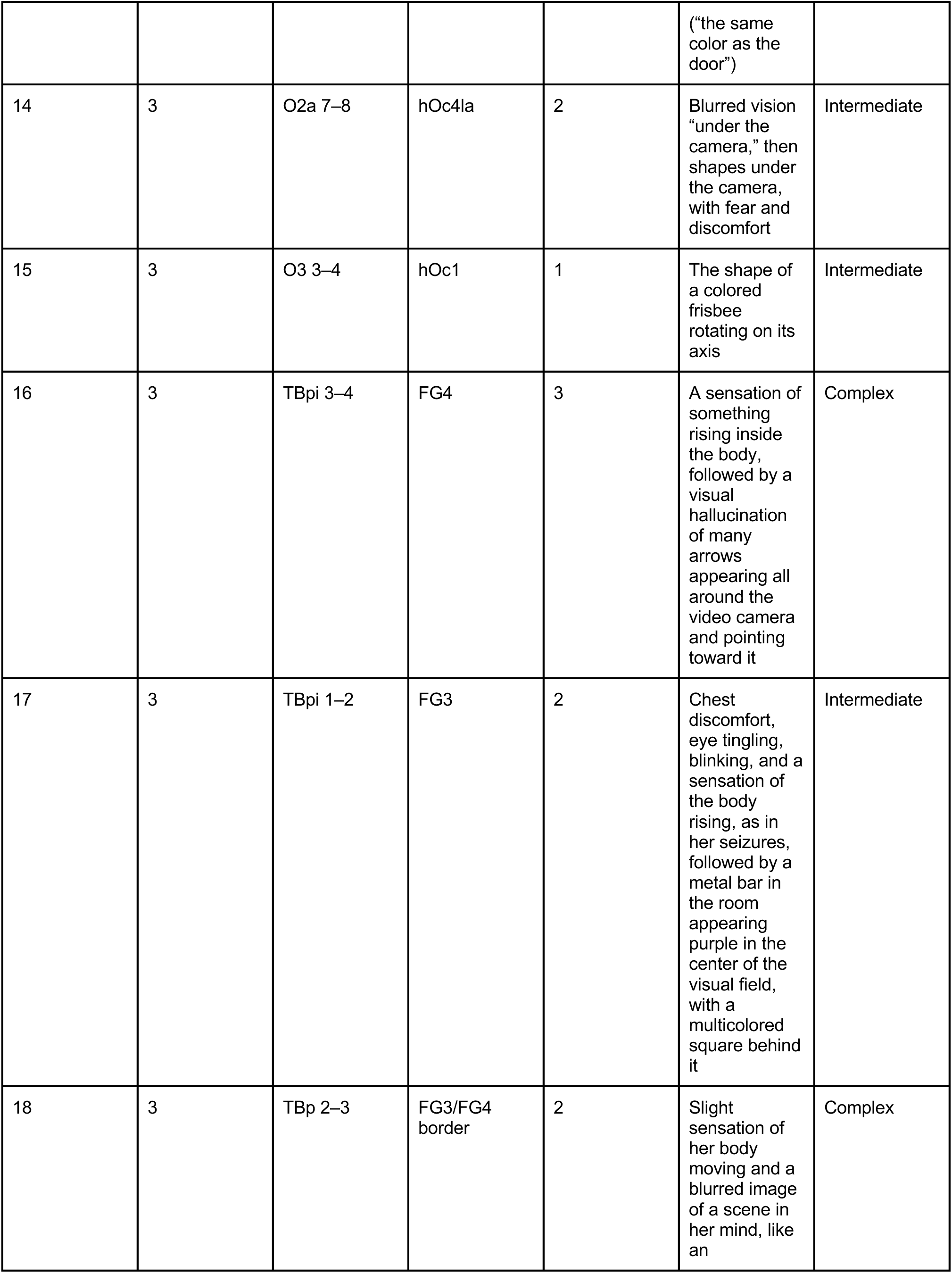
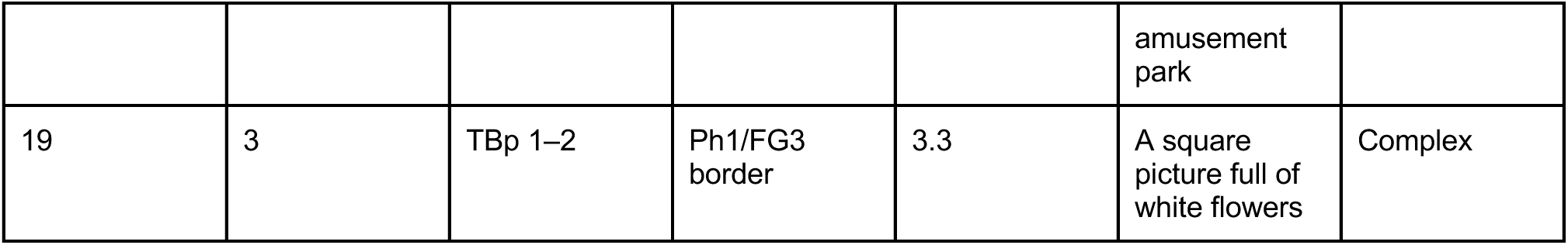
Visual experiences evoked at each stimulation site, numbered as in Figure 1. All stimulations were bipolar, at 50 Hz; intensity is the lowest that evoked the reported experience. FG1, FG3 and FG4, fusiform gyrus areas 1, 3 and 4; hIP7, human intraparietal area 7; hOc1, human occipital area 1 (primary visual cortex); hOc2, human occipital area 2; hOc4d, hOc4la and hOc4v, dorsal, lateral-anterior and ventral subdivisions of human occipital area 4; mA, milliampere; Ph1, parahippocampal area 1.

The analysis included every contact pair whose stimulation evoked a visual experience, regardless of location, subject only to the following exclusions. Stimulations involving white-matter contacts or contacts adjacent to the optic radiation, or eliciting pathological sEEG activity, including local or propagating afterdischarges [13] and clinical or subclinical ictal discharges, were excluded from the analysis. Only Patient 1 had left-hemisphere electrodes; none of his left-hemisphere stimulations met the inclusion criteria (Table S1). When increasing the intensity at a given contact pair elicited different manifestations, only the response obtained at the lowest intensity that evoked a classifiable visual manifestation was retained to limit recruitment of structures beyond the stimulation site. The same rule applied when the lowest effective intensity elicited an illusion. Consequently, contact pair Tppi 3–4 (Patient 3) was excluded even though stimulation at 3 mA evoked colored spots (Table S1). If stimulation provoked a seizure, DIECS was suspended until full clinical recovery and restoration of the usual sEEG background.

### Classification of stimulation-induced visual experiences

Immediately after each stimulation train, patients provided a verbal description of the perceived phenomenon. A visual manifestation was included in the primary analysis only if the patient’s description clearly identified it and provided sufficient detail to allow classification according to the scheme proposed by Lee et al. [4]. Simple-form responses were unstructured visual phenomena, such as white, black, or colored spots, blobs, or flashes of light. Intermediate-form responses comprised elementary geometric forms, including triangles, diamond-like shapes, and stars, which could be achromatic or colored and either stationary or moving. The boundary between simple and intermediate responses is not sharp. For example, we classified the oval luminous figure at site 3 as simple, but reclassifying it as intermediate would leave the pattern unchanged (8 simple, 4 intermediate, and 0 complex responses at the 12 occipital or intraparietal sites). Complex responses consisted of formed and integrated visual experiences, such as people, objects, pictures, or scenes. Because our main interest concerned visual experiences arising without a corresponding external stimulus, as in visual mental imagery, the primary analysis included only responses that evoked new visual content, whether experienced internally or in the external scene. Responses that only altered the perception of the existing scene, such as apparent movement, oscillation, or blurring of the visual field, selective clarity of attended objects, or a color change of a real object, were classified separately as illusions, following Lee et al., and are listed in Table S1. When stimulation both evoked new visual content and altered the appearance of the existing scene, the response was classified as simple, intermediate, or complex according to the newly evoked content. Three authors (V.F., A.C., and D.H.) classified the reports before the Jülich-Brain labels of the stimulation sites were available, and the classification was then reviewed and discussed with two other authors (S.D.S. and P.B.).

### Anatomical localization of bipolar stimulation sites

Bipolar stimulation sites were anatomically localized with the epiSTIM software platform (STIM-IMA SEEG toolbox) [14]. Electrode contacts were reconstructed from postoperative CT images coregistered to the preoperative T1-weighted MRI and localized in each patient’s native space and MNI152 standard space. Each bipolar stimulation site was defined as the three-dimensional midpoint between the two stimulated contacts. The midpoint’s MNI coordinates were used for anatomical annotation (Table S2). Both contacts of every pair included in the primary analysis were verified to lie in cortical gray matter in the patient’s native space. When the two contacts lay on opposite banks of a sulcus, the midpoint could fall outside cortical gray matter, and the stimulation was considered to involve both cortical banks. Sites were labeled using the Jülich-Brain probabilistic cytoarchitectonic atlas (version 3.1) [15], including the cytoarchitectonic subdivisions of the fusiform gyrus described by Caspers et al. [10] and Lorenz et al. [9], with FG1 and FG2 on the medial and lateral posterior fusiform gyrus and FG3 and FG4 on the medial and lateral mid-fusiform gyrus, as implemented in epiSTIM. The area label of each site is the Jülich-Brain area assigned to the midpoint. Sites 18 and 19 lay at the border between two areas. Descriptions of contact location in gyral terms, such as lateral occipital cortex or cuneus, refer to each patient’s own anatomy and parcellate the cortex differently from the cytoarchitectonic atlas, so the two can name the same site differently (for example, site 1 lay in the lateral occipital gyrus and in area hIP7). For visualization and quality control, all stimulation-site midpoints and the Jülich-Brain parcellation were displayed in MNI space in an interactive 3D Slicer scene.

### Ethics

All participants provided written informed consent to participate in the study (INSERM projects C11-16 and C19-55, approved by the local ethics committee, CPP Paris VI). All procedures followed the relevant ethical regulations and guidelines. The study was not preregistered. The authors used Claude (Anthropic) and ChatGPT (OpenAI) to edit the text for grammatical correctness and clarity and to draft the figure legend and Tables 1, S1, and S2; the authors reviewed and revised these passages and take full responsibility for the content.

### Statistical Analyses

The analysis is descriptive, and no statistical test was applied. No statistical method was used to predetermine sample size, which was set by the number of patients meeting the inclusion criteria stated above. No data were excluded beyond the exclusions specified above. There was no group allocation, so randomization did not apply. Patients were not told the onset of each stimulation train. The initial classification of reports preceded the assignment of Jülich-Brain labels (see Classification of stimulation-induced visual experiences).

### Code availability

No custom code was developed for this study. Stimulation sites were localized with the epiSTIM platform described above.

## Results

Table 1 and Figure 1 summarize the 19 sites included in the primary analysis. Of these, 12 lay in occipital or adjacent intraparietal cortex (9 simple, 3 intermediate, 0 complex) and 7 in or at the border of fusiform areas (0 simple, 3 intermediate, 4 complex). Because several sites come from the same patient, sites are not independent observations, and we report these counts descriptively. Excluding site 1, the only intraparietal site, leaves the pattern unchanged (11 occipital sites: 8 simple, 3 intermediate, 0 complex). Among the four sites with complex responses, Table S1 documents a similar response at higher intensity at site 16; at sites 18 and 19, higher intensity evoked nonvisual manifestations; site 6 was not retested. The MNI coordinates and Jülich-Brain labels of these sites are given in Table S2. Table S1 lists all other stimulation-evoked manifestations observed in the three patients, including nonvisual manifestations, responses elicited by repeat stimulation, visual illusions, and visual experiences that did not meet the inclusion criteria for the primary analysis.

**Figure 1.**
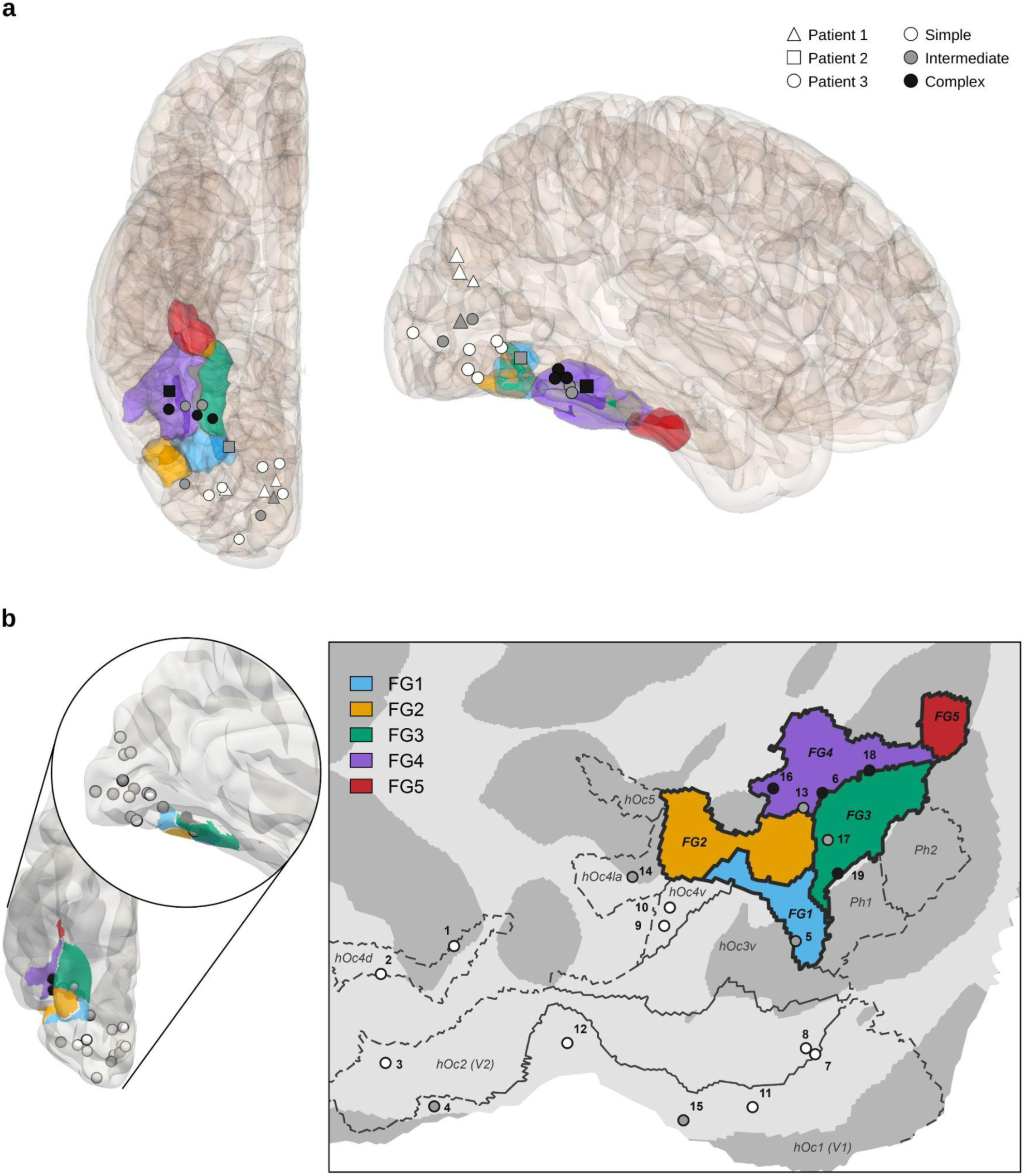
Anatomical location and phenomenology of stimulation-induced visual experiences. a, Bipolar stimulation sites (midpoints between the two stimulated contacts) on a semi-transparent rendering of the right hemisphere in MNI space, in ventral (left) and lateral (right) views, with the fusiform areas FG1 (blue), FG2 (orange), FG3 (green), FG4 (purple), and FG5 (red) of the Jülich-Brain atlas. Marker color indicates the class of the evoked visual experience (white, simple; gray, intermediate; black, complex), and marker shapes indicate the patients (triangles, Patient 1; squares, Patient 2; circles, Patient 3). b, The same sites in MNI space, with the Jülich-Brain cytoarchitectonic parcellation (version 3.1). Left, ventral and lateral (inset) views of a three-dimensional volume rendering of the right occipito-temporal lobe, with the fusiform areas colored as in a. Right, flattened map of the right ventral occipito-temporal cortex, with the borders of the Jülich-Brain areas in dark black lines. Markers indicate the class of the evoked visual experience, coded as in a. On the flattened map, numbers refer to the stimulation sites listed in Table 1. Projection onto the flattened map can displace individual sites relative to their position in the volume rendering shown at left.

### Patient 1

Electrical stimulation in Patient 1 elicited only simple or intermediate visual experiences (Table 1 and Figure 1), all from posterior cortical sites. Stimulation of intraparietal area hIP7 (site 1) at 50 Hz and 2 mA, with contacts in the right lateral occipital cortex, produced something flickering in the left visual hemifield. Stimulation of dorsal occipital area 4 (hOc4d; site 2) at 50 Hz and 2 mA induced the repeated appearance and disappearance of an unspecified luminous percept. Stimulation of occipital area V2 (hOc2; site 3) [16] at 50 Hz and 2 mA, with contacts in the right cuneus, elicited an oval luminous figure in the center of the visual field associated with flickering of the visual field. Stimulation of primary visual cortex (hOc1; site 4) at 50 Hz and 4 mA induced the perception of two colorless luminous circles in the left visual hemifield. No complex visual experiences were reported, even when intensity was increased at sites 2–4 or when additional occipital and temporal sites were stimulated (Table S1).

### Patient 2

Stimulation of several sampled sites in Patient 2 elicited visual experiences (Table 1 and Figure 1). Stimulation of the right posterior fusiform area FG1 (site 5) at 50 Hz and 1 mA elicited an intermediate experience, described as a spiral descending from the upper-left visual hemifield. By contrast, stimulation of the right lateral mid-fusiform area FG4 (site 6) at 50 Hz and 2 mA elicited three blurred images of people that the patient could not describe in detail. The only complex visual experience reported by this patient thus arose from the more anterior fusiform subdivision, FG4. Stimulation of the right intraparietal cortex (hIP4) and of the adjacent temporoparietal cortex at 1 mA instead altered the perception of the existing scene, which appeared to move or oscillate vertically; these illusions are listed in Table S1.

### Patient 3

Electrical stimulation in Patient 3 elicited visual experiences from right occipital and fusiform sites and from a site at the Ph1/FG3 border (Table 1 and Figure 1). Sites 18 and 19 were stimulated during the first sEEG exploration and the remaining sites during the second; complex visual experiences arose in both explorations. At the FG3/FG4 border (site 18; 2 mA), stimulation produced a blurred image resembling an amusement park, experienced “in her mind,” together with a sensation of body movement. Stimulation at the Ph1/FG3 border (site 19; 3.3 mA) produced an image resembling a square picture filled with white flowers. Stimulation of FG4 at site 16 (3 mA) evoked a rising bodily sensation followed by a complex visual experience of many arrows appearing all around the video camera and pointing toward it. We classified this response as complex because the arrows formed a spatial configuration around the camera rather than a single geometric form; classifying it as intermediate would leave the pattern intact, with complex responses at three of the seven fusiform sites and at none of the 12 occipital or intraparietal sites. Stimulation of FG3 (site 17; 2 mA) elicited bodily and ocular sensations that she recognized from her seizures, followed by a multicolored square behind a metal bar in the room. The bar itself appeared purple, an illusory change of a real object. The response was classified as intermediate on the basis of the newly seen square. FG4 stimulation at site 13 (1.5 mA) also elicited an intermediate experience, a small empty square in the center of the visual field. By contrast, stimulation of the occipital sites (sites 7–12, 14, and 15) elicited flashes, colored spots or lights, or simple shapes, occasionally accompanied by fear or discomfort. No complex visual experience was reported after occipital stimulation. Blurred vision after stimulation of the lateral occipital area hOc4la and of the parahippocampal area Ph1 was classified as an illusion (Table S1).

## Discussion

In this anatomically detailed multiple single-case series, direct stimulation elicited complex visual experiences from sites in or adjacent to the right mid-fusiform areas FG3 and FG4. By contrast, the sampled occipital sites and the posterior fusiform FG1 site elicited only simple or intermediate phenomena, such as flickering lights or geometric shapes. Occipital stimulation never elicited a complex experience, and mid-fusiform stimulation never a simple one. Only intermediate forms arose from both regions. This distribution supports a mid-fusiform contribution to complex visual content.

This anatomical pattern confirms and refines the observations of Penfield and Perot [3], who reported predominantly elementary visual experiences after occipital stimulation and complex visual experiences only after temporal stimulation, mostly of the right hemisphere and including sites on the basal temporal surface. Salanova et al. [17] found that intraoperative occipital stimulation reproduced habitual auras in 37% of 29 patients; all were elementary visual hallucinations. In two patients, right posterior temporal stimulation also evoked complex visual experiences. They suggested that complex hallucinations during occipital seizures could reflect spread to temporal structures. Our observations also extend previous stimulation studies [4, 6, 18, 19] by associating differences in subjective experience with cytoarchitectonic subdivisions of the fusiform gyrus. In the series by Yuan et al. [19], comprising 30,404 stimulated sites in 335 patients, memory-related visual experiences, which differ from the complexity classes used here, were elicited at 30 of 1,496 fusiform sites, in 17 of 195 patients. After correction for multiple comparisons, only the fusiform response rate significantly exceeded that of the superior temporal sulcus, their reference region; occipital and parieto-occipital responses were rare.

Two patient series have nonetheless reported complex visual hallucinations after posterior stimulation. Andelman-Gur et al. [20] used 1,165 subdural contacts spaced 10 mm apart and 134 depth contacts spaced 5 mm apart, with stimulation currents of 1–10 mA and 1–4 mA, respectively. They assigned sites to lobes and gyri. Complex hallucinations also followed frontal stimulation, as previously described for the left prefrontal cortex [21]. Andelman-Gur et al. found that complex visual hallucinations clustered in the left occipital lobe but were widely dispersed in the right hemisphere. They suggested that this difference might reflect hemispheric asymmetries in white-matter organization. In our patients, increasing the intensity at the same occipital contact pairs still produced only simple or intermediate experiences. Intensities reached 4 mA at site 2, 3 mA at site 3, and 8 mA at site 4 in Patient 1, and 2.5 mA in Patient 3 (Table S1). By contrast, mid-fusiform stimulation at 2–3.3 mA produced complex visual experiences. At FG4 site 13, repeated stimulation at 3 mA evoked a square containing words and phrases the patient could not report, a gain in content that no occipital pair showed (Table S1). Jonas et al. [6] evoked face hallucinations with depth electrodes, mostly in the right cuneus and parieto-occipital sulcus, but included responses accompanied by afterdischarges limited to the immediate vicinity of the stimulated site. Neither study assigned sites to cytoarchitectonic areas, so it is not possible to adjudicate whether their posterior responses originated in early visual cortex or in adjacent higher-order cortex. The frontal responses, and possibly the cuneus and parieto-occipital sulcus responses, suggest that complex visual experiences can be evoked from nodes of the underlying networks that are distant from the ventral temporal cortex.

The content of the evoked experiences may also reflect the selectivity of the stimulated cortex. Stimulation of the right fusiform face-selective cortex can evoke face-related hallucinations [22] or distortions of perceived faces [23], and the change in a real object’s perceived color that we elicited by stimulating FG3 (site 17 in Table 1) resembles color phenomena previously reported after stimulation of a color-biased medial fusiform region [5].

Fusiform organization may relate not only to content complexity but also to whether evoked experiences seem internal or external. Yuan et al. [19] also reported that anterior fusiform stimulation more often evoked internally oriented reminiscences, whereas more posterior fusiform stimulation more often elicited experiences perceived as occurring in external space. Although Yuan et al.’s anatomical subdivisions do not map directly onto FG1–FG4, their findings raise the possibility that fusiform organization contributes not only to visual content but also to whether that content is experienced as internally or externally generated. The possible role of fusiform cortex in attributing visual experiences to internal or external sources is also relevant to reality monitoring. Using fMRI, Dijkstra et al. [24] found that bilateral fusiform activity tracked the combined strength of imagined and perceived signals, with left fusiform activity predicting when an imagined pattern was judged externally present. Fusiform activity was also coupled to the anterior insula, where activity tracked categorical judgments of stimulus presence. Dijkstra et al. related their findings to the consistent involvement of the fusiform gyrus in visual mental imagery. Their left fusiform cluster lay near the Fusiform Imagery Node (FIN), a region within cytoarchitectonic area FG4 that is consistently active during imagery [25–27].

These findings bear on theories of voluntary visual mental imagery, although they do not directly test them. As mentioned in the introduction, some current models propose that mental images acquire their visual quality in early visual cortex [1]. Our data show that complex visual experiences can be initiated from high-level ventral temporal cortex. Among the evoked phenomena, these complex experiences come closest to typical visual mental images, which usually represent objects, people, or scenes; at site 18, the patient described the image as being “in her mind.” They differ from voluntary imagery in arising without intention. Occipital stimulation itself evoked vivid, if elementary, visual phenomena, so our data alone remain compatible with a hierarchical account; the contrast between sites concerns the complexity of content rather than visual quality as such. However, converging evidence for a dissociation between occipital and ventral temporal cortex comes from lesion neuropsychology, epilepsy semiology, and neuroimaging. After brain damage, vivid and detailed imagery may survive substantial occipital injury [2, 28], whereas temporal lesions can impair it [29, 30]. In epileptic seizures, elementary visual auras are commonly associated with occipital seizure foci [31], whereas complex scenes and experiential phenomena are more closely linked to temporal and occipito-temporal networks [32, 33]. Neuroimaging also implicates high-level ventral temporal and frontoparietal regions in imagery, with less consistent involvement of early visual cortex [25, 26]. Together, the lesion and neuroimaging findings challenge the necessity of early visual cortex for imagery and motivate a network-level account [34]. Our stimulation results are compatible with this account, although they do not show that voluntary imagery and stimulation-evoked hallucinations rely on the same mechanisms.

One possible difference concerns hemispheric laterality. Neuroimaging and lesion evidence implicates the left-hemisphere FIN in voluntary imagery [25, 26, 35, 36]. By contrast, complex visual experiences in this study followed stimulation of right mid-fusiform sites, consistent with earlier evidence of a right-hemisphere bias in stimulation-evoked visual phenomena [3, 6]. This difference may reflect distinct mechanisms of voluntary imagery and externally induced experiences [37]. Network control theory offers a testable mechanistic hypothesis for how such a hemispheric asymmetry could arise. Network control analyses of structural connectomes predict greater right-to-left than left-to-right control for most brain systems [38]. We hypothesize that right mid-fusiform stimulation could alter left FIN activity through interhemispheric connections [39–41]. Altered FIN activity could then contribute to imagery-like experiences outside voluntary control. Because only Patient 1 had left-hemisphere electrodes, and none of his stimulations evoked a complex response, the right-sided localization may partly result from electrode sampling, although the right-hemisphere bias reported in stimulation studies since Penfield and Perot [3] makes sampling unlikely to be the sole explanation. A direct test would combine right fusiform stimulation with concurrent recording from the left FIN.

More broadly, interactions between low- and high-level visual cortex and frontoparietal networks, which heterarchical models of imagery emphasize, may influence whether visual content becomes consciously accessible [34, 42–45]. In a patient with cortical blindness, Hauw et al. [46] found that visual stimuli still activated partially preserved V1, while functional connectivity between V1 and prefrontal cortex was reduced. This case suggests that communication between early visual and prefrontal regions may be critical to conscious visual perception, consistent with an early proposal [47]. For voluntary imagery, findings in congenital aphantasia are also consistent with the importance of communication between temporal and frontal regions [26, 36, 48–50].

Several limitations qualify these findings. First, there were only three patients, and electrode implantation was determined by clinical needs. Only Patient 3 contributed both occipital and fusiform sites. In addition, inclusion required stimulation-evoked responses from at least two phenomenological classes. We therefore cannot estimate the probability of complex responses across the full occipito-temporal cortex, exclude unsampled occipital sites that might evoke complex visual experiences, or establish a continuous gradient of response complexity. Second, the patients had long-standing drug-resistant epilepsy, and functional reorganization cannot be ruled out. Moreover, complex responses arose in or near the epileptogenic zones of Patients 2 and 3, and the response at site 17 began with sensations that the patient recognized from her seizures. Patient 3, who contributed three of the four complex responses, had habitual seizures beginning with complex visual hallucinations, so some responses may partly reflect activation of her epileptic network. However, in the large series of Yuan et al. [19], in which all effective sites lay outside the epileptogenic zone, fusiform stimulation also evoked memory-related visual experiences. Third, bipolar stimulation is spatially constrained but not exclusively local; physiological activation of connected areas, including occipital cortex, may occur without detectable afterdischarges. Fourth, descriptions and classifications of subjective experiences involve interpretive judgments. Some reports combined visual, bodily, and affective phenomena, and the distinction between intermediate and complex forms is not always sharp, nor is the distinction between new visual content and an altered perception of the existing scene, which rests on brief verbal reports.

Despite these limitations, direct stimulation in three patients adds cytoarchitectonically localized causal evidence that perturbing right mid-fusiform cortex, but not the sampled occipital sites, can trigger complex visual experiences. These observations are compatible with lesion and neuroimaging evidence challenging the view that mental images acquire their visual quality in early visual cortex, and identify the mid-fusiform gyrus and its interhemispheric and frontoparietal connections as promising targets for future causal studies.

## Data availability

Site-level data supporting the findings, including anatomical labels, MNI coordinates, stimulation parameters, and reported experiences, are provided in Table 1 and Tables S1 and S2. No dataset was deposited in a public repository. The raw sEEG recordings and the clinical records are identifiable patient data and cannot be made publicly available. De-identified data are available from V.F. on reasonable request, within the limits set by the patients’ consent and French regulations on health data.

## Acknowledgments

We thank the patients for their participation. The work of P.B. is supported by the Agence Nationale de la Recherche through ANR-16-CE37-0005 and ANR-10-IAIHU-06, by the Fondation pour la Recherche sur les AVC through FR-AVC-017, and by the Paris Brain Institute grant ViBER - Vision Beyond External Reality.

## Author contributions

D.H. and V.F. contributed equally. P.B. conceived the study. V.F. and V.N. were responsible for the sEEG investigations and stimulation sessions. V.F., A.C., and D.H. classified the visual reports, and S.D.S. and P.B. reviewed the classification. V.F., S.F.V., and K.L. localized the stimulation sites. J.L. and V.F. prepared Figure 1. P.B., D.H., V.F., and S.D.S. wrote the manuscript with input from all authors. All authors approved the final version.

## Supplementary Material

**Table S1:**
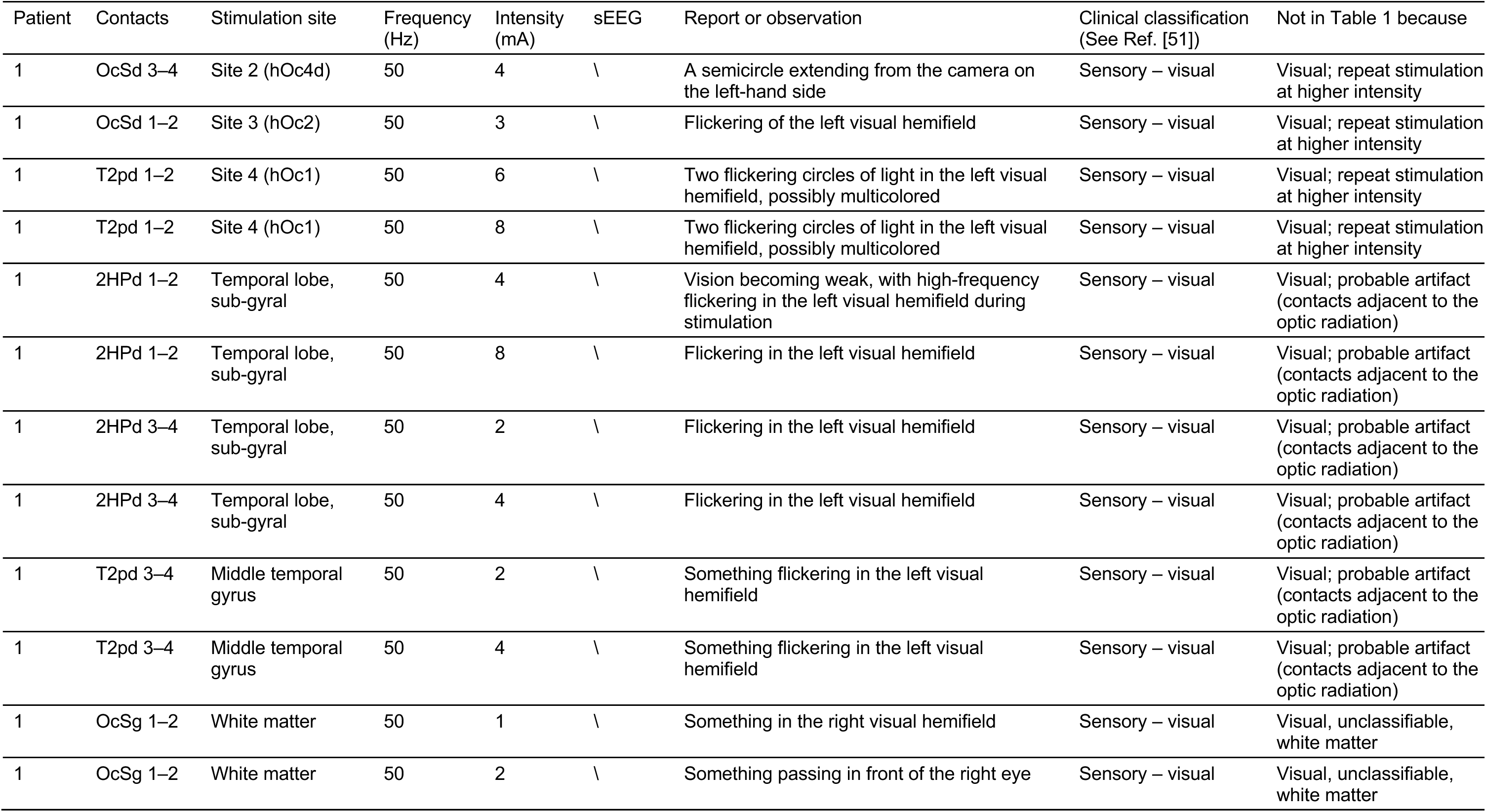
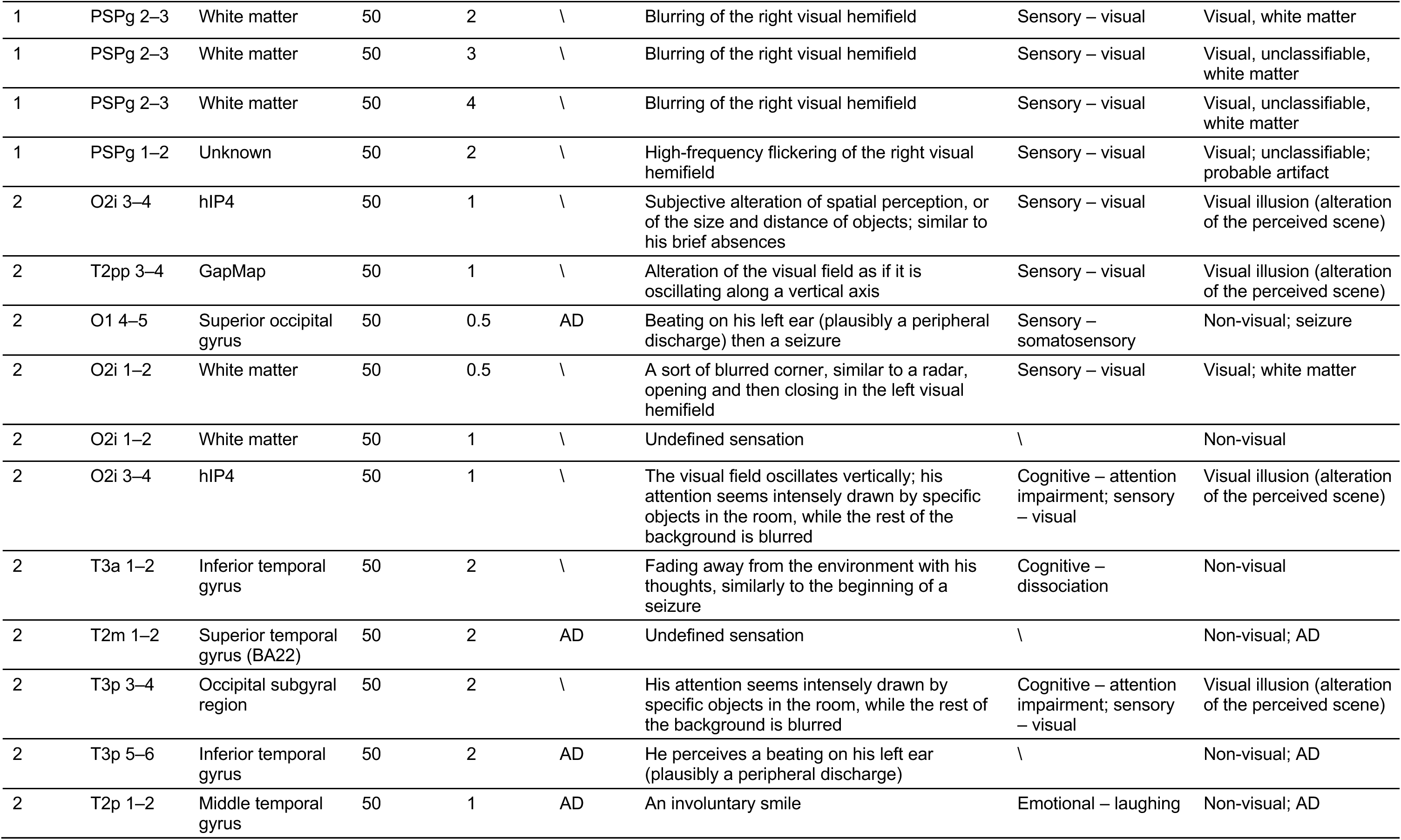
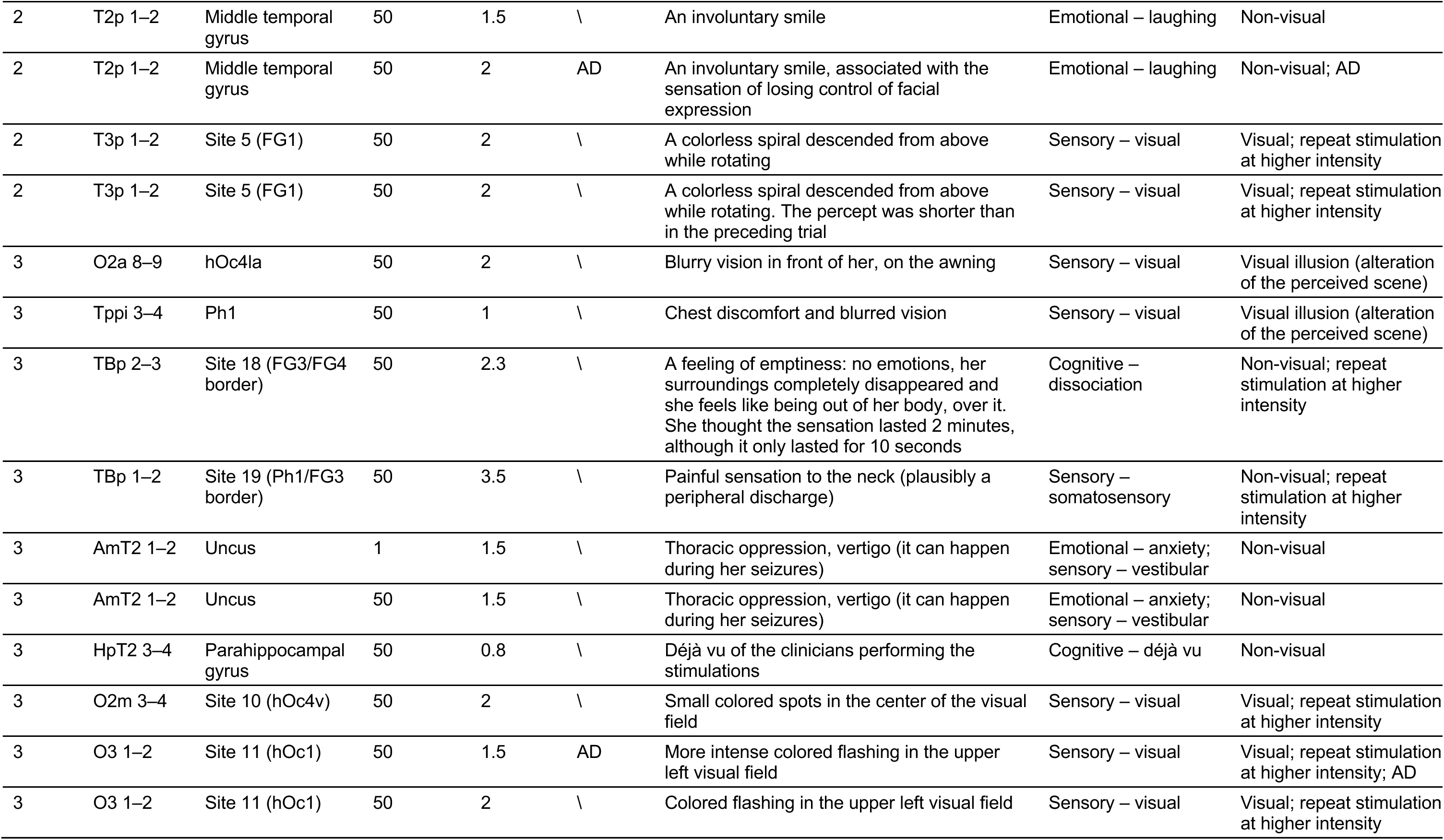
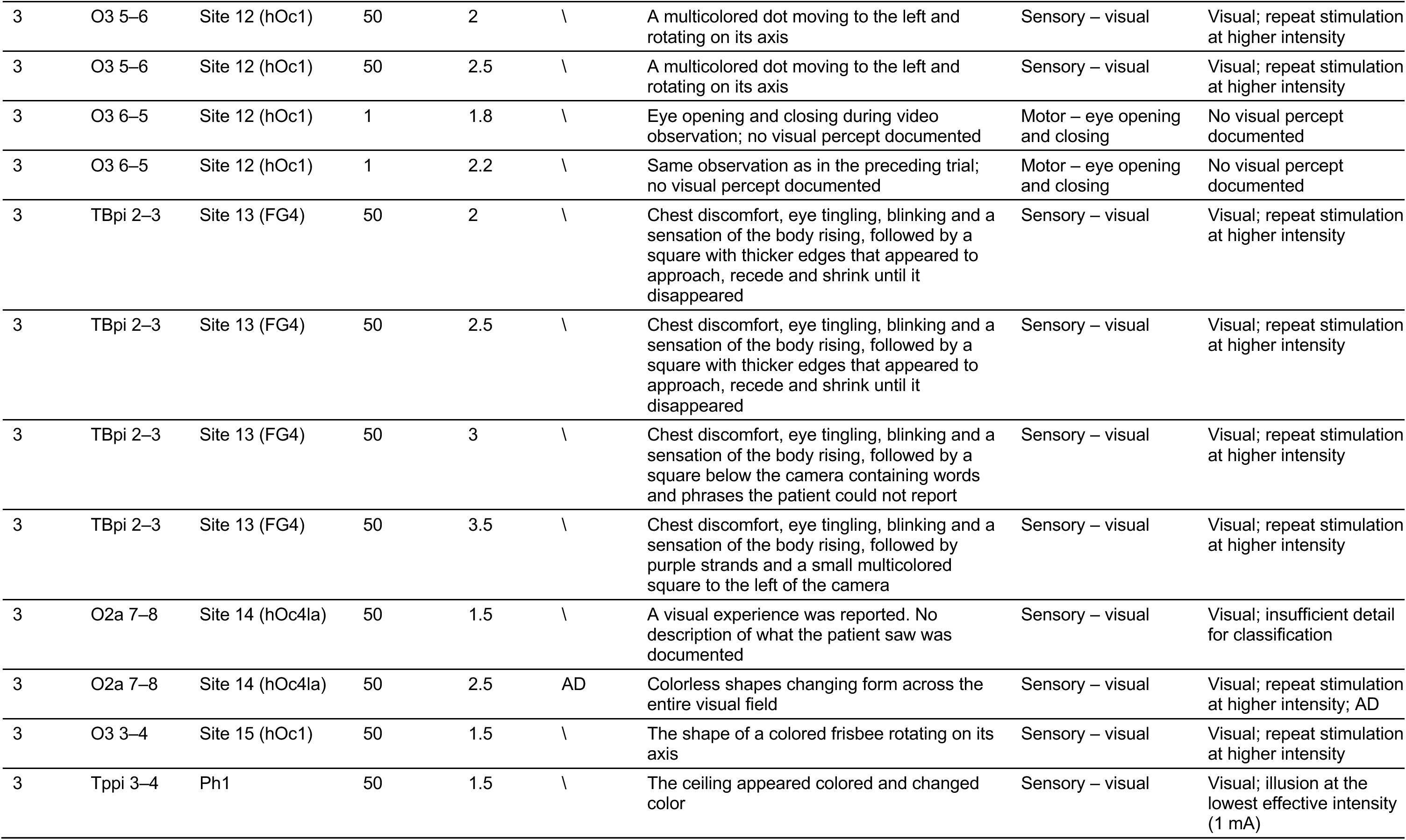
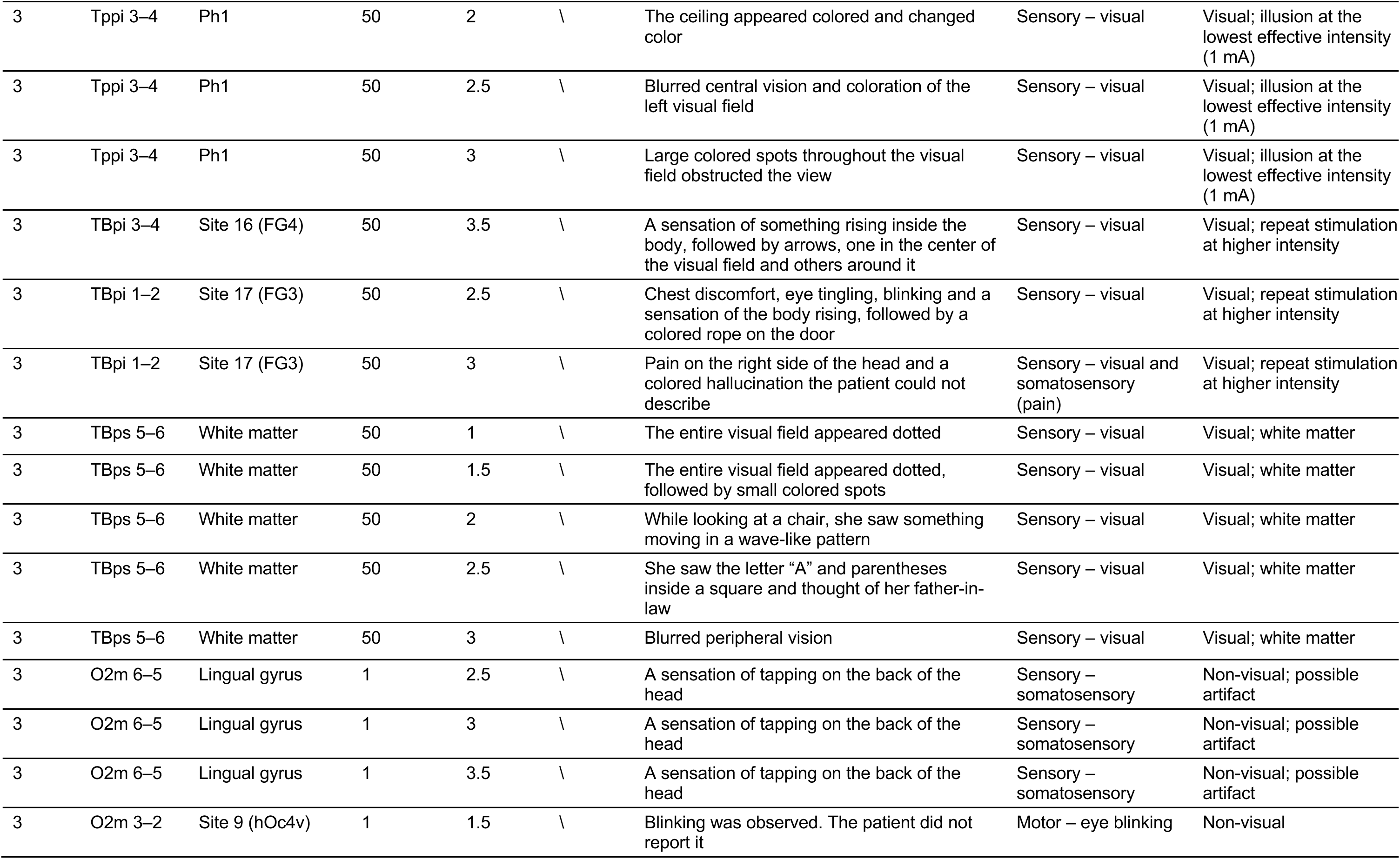
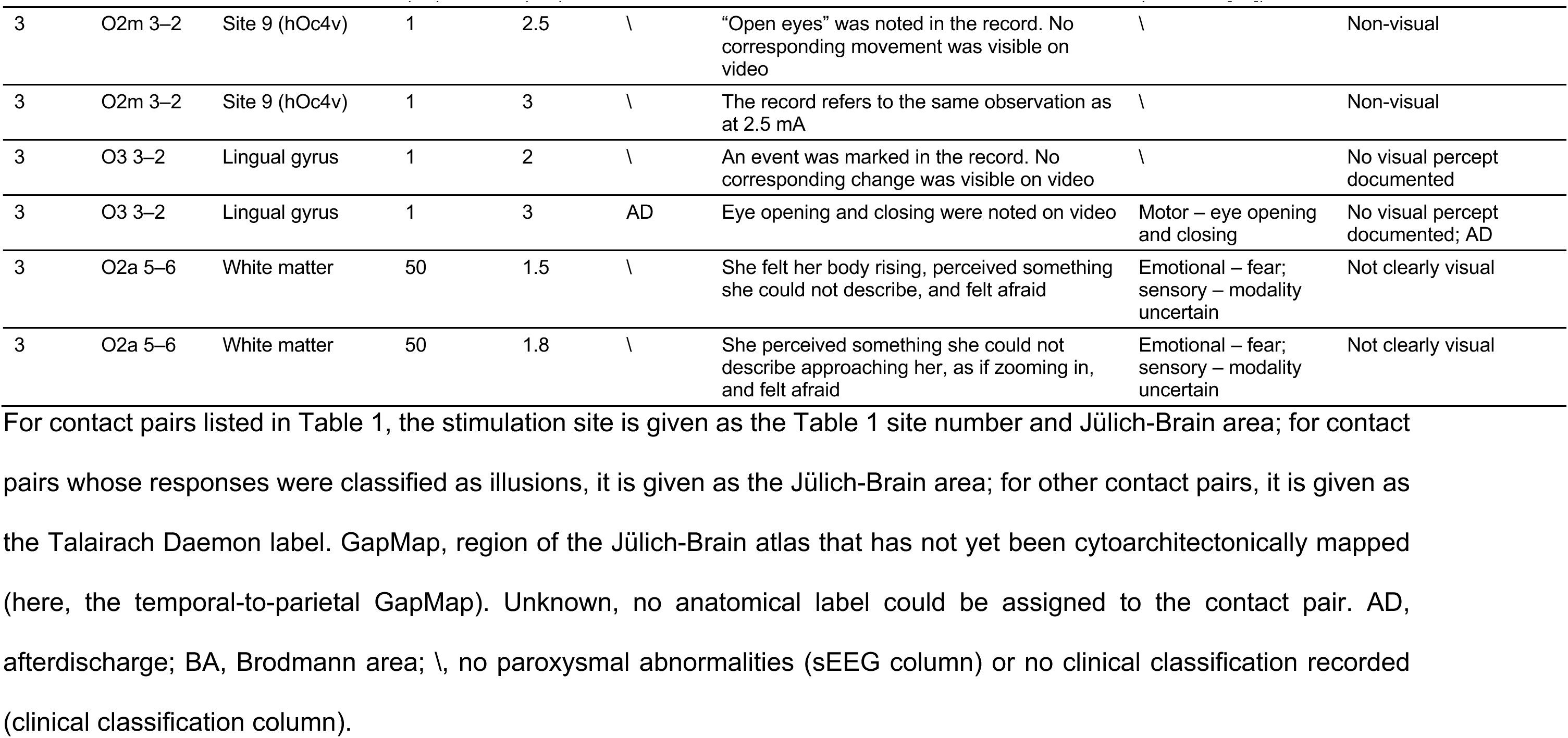
Stimulation-induced manifestations not included in Table 1.

**Table S2:** MNI coordinates and Jülich-Brain areas of the stimulation sites.

| Site | Patient | Contacts | MNI x (mm) | MNI y (mm) | MNI z (mm) | Jülich-Brain area |
|---|---|---|---|---|---|---|
| 1 | 1 | T2pd 5–6 | 27.3 | −82.5 | 18.8 | hIP7 |
| 2 | 1 | OcSd 3–4 | 13.6 | −84.7 | 27.3 | hOc4d |
| 3 | 1 | OcSd 1–2 | 9.6 | −79.8 | 20.0 | hOc2 |
| 4 | 1 | T2pd 1–2 | 10.3 | −83.5 | 6.8 | hOc1 |
| 5 | 2 | T3p 1–2 | 25.4 | −64.0 | −6.3 | FG1 |
| 6 | 2 | T3m 1–2 | 44.3 | −44.6 | −17.5 | FG4 |
| 7 | 3 | O2a 1–2 | 8.5 | −70.4 | −0.1 | hOc2 |
| 8 | 3 | O2a 2–3 | 14.1 | −71.5 | 0.5 | hOc2 |
| 9 | 3 | O2m 2–3 | 27.0 | −76.6 | −12.5 | hOc4v |
| 10 | 3 | O2m 3–4 | 31.7 | −79.2 | −10.9 | hOc4v |
| 11 | 3 | O3 1–2 | 7.3 | −80.4 | −0.9 | hOc1 |
| 12 | 3 | O3 5–6 | 22.3 | −96.3 | 1.5 | hOc1 |
| 13 | 3 | TBpi 2–3 | 39.0 | −49.5 | −16.6 | FG4 |
| 14 | 3 | O2a 7–8 | 40.6 | −77.6 | 2.3 | hOc4la |
| 15 | 3 | O3 3–4 | 14.7 | −88.0 | 0.3 | hOc1 |
| 16 | 3 | TBpi 3–4 | 44.5 | −50.5 | −15.2 | FG4 |
| 17 | 3 | TBpi 1–2 | 33.5 | −48.9 | −17.9 | FG3 |
| 18 | 3 | TBp 2–3 | 35.6 | −52.8 | −11.5 | FG3/FG4 border |
| 19 | 3 | TBp 1–2 | 30.4 | −53.7 | −13.0 | Ph1/FG3 border |

Coordinates are those of the midpoint between the two stimulated contacts, in MNI152 space. Areas are as in Table 1; at sites 18 and 19 the midpoint lay at the border between two areas.

